# Decomposing Cognitive Processes in the mPFC During Self-Thinking

**DOI:** 10.1101/2024.11.13.623526

**Authors:** Marie Levorsen, Ryuta Aoki, Constantine Sedikides, Keise Izuma

**Affiliations:** School of Psychology, University of Southampton, Southampton, SO17 1BJ, UK; Graduate School of Humanities, Tokyo Metropolitan University, Tokyo, 192-0397, Japan; Human Brain Research Center, Graduate School of Medicine, Kyoto University, Kyoto, 606-8501, Japan; School of Economics & Management, Kochi University of Technology, Kochi, 780-8515, Japan; Research Institute for Future Design, Kochi University of Technology, Kochi, 780-8515, Japan

**Keywords:** self, self-reference, introspection, memory, default-mode, variance partitioning, multivariate pattern regression, multi-voxel pattern analysis, representational similarity analysis, medial prefrontal cortex

## Abstract

Past cognitive neuroscience research has demonstrated that thinking about both the self and other activate the medial prefrontal cortex (mPFC), a central hub of the default mode network. The mPFC is also implicated in other cognitive processes, such as introspection and autobiographical memory, rendering elusive its exact role during thinking about the self. Specifically, it is unclear whether the same cognitive process explains the common mPFC involvement or distinct processes are responsible for the mPFC activation overlap. In this preregistered functional magnetic resonance imaging study, we investigated whether and to what extent mPFC activation patterns during self-reference judgment could be explained by activation patterns during the tasks of other-reference judgment, introspection, and autobiographical memory. Multi-voxel pattern analysis showed that only in the mPFC were neural responses both concurrently different and similar across tasks. Furthermore, multiple regression and variance partitioning analyses indicated that each task (i.e., other-reference, introspection, memory) uniquely and jointly explained significant variance in mPFC activation during self-reference. These findings suggest that the self-reference task involves multiple cognitive processes shared with other tasks, and the mPFC is the unique place where necessary information is gathered and integrated for judgments based on internally constructed representations.

## Introduction

Thinking about the self and expressing who one is to others are fundamental aspects of human experience. The self has fascinated researchers for more than a century (James, 1890; Cooley, 1902; Sedikides and Gregg, 2003; Baumeister, 2023). Reflecting this enduring interest, the intricate neural architecture of the self has been a persistent focus of inquiry (Wagner et al., 2019; Frewen et al., 2020). Using neuroimaging methods such as functional magnetic resonance imaging (fMRI), studies have established that the midline structures, the medial prefrontal cortex (mPFC) and posterior cingulate cortex (PCC), are active when individuals judge if a presented personality trait or attitudinal statement describes them (Johnson et al., 2002; Kelley et al., 2002; Izuma et al., 2010; for meta-analyses, see Denny et al., 2012; Murray et al., 2012).

Although the robust link between the mPFC and self-reference processing raised the possibility that the primary function of the mPFC is involvement in self-relevant information (Kelley et al., 2002; Northoff, 2016), the mPFC is also involved in thinking about other people (Denny et al., 2012; Murray et al., 2012). Previous fMRI studies have observed an overlap in mPFC activation between the self-reference task and other-reference task (judging if an adjective describes a close other; (Ochsner et al., 2005; Vanderwal et al., 2008; Krienen et al., 2010; see also a recent human electrocorticography [ECoG] study [Tan et al., 2022] showing high similarity in neural responses between self-vs. other-reference processing across the whole brain). Based on these observations, some researchers (Gillihan and Farah, 2005; Legrand and Ruby, 2009) criticized the self-specific view of the mPFC, arguing that certain general cognitive processes are common to self- and other-reference processing. For example, inferential processing and memory recall seems to be common to both (Legrand and Ruby, 2009). In other words, the mPFC activation during the self-reference task might not be related to the self specifically, but rather it is a result of general cognitive processes that take place during the self-reference task, as well as during other tasks. Indeed, the mPFC and PCC are also known to be activated by autobiographical memory (Kim, 2012; Martinelli et al., 2013) and by decision-making based on internal or subjective criteria (e.g., moral decision-making; Nakao et al., 2012).

From a broader perspective, the mPFC and PCC are considered the core hubs of the default mode network – a network of brain regions that show heightened activation at rest (Andrews-Hanna et al., 2010; Andrews-Hanna, 2012). These regions are activated by a variety of tasks that depend on internally constructed representations (i.e., self-generated mental contents), including not only self- and other-reference processing, as well as autobiographical memory, but also introspection (thinking about one’s own emotional states), episodic future thinking, creativity, affective decision-making, and spatial navigation (Andrews-Hanna et al., 2014; Buckner and DiNicola, 2019; Wen et al., 2020; Menon, 2023). For the past two decades, researchers have attempted to identify a key common cognitive process that explains mPFC’s involvement in these distinct tasks, such as self-projection (Buckner and Carroll, 2007), self-relevant processing (D’Argembeau et al., 2010; Northoff, 2016), scene construction (Hassabis and Maguire, 2007), affective meaning generation (Roy et al., 2012), and internal narrative construction (Menon, 2023). However, these attempts are often based on univariate activation overlap or meta-analyses (Spreng et al., 2009; Lieberman et al., 2019; Menon, 2023). It is increasingly recognized that univariate activation overlap does not constitute strong evidence for a common cognitive process across tasks (Woo et al., 2014; Levorsen et al., 2023). Thus, experimental evidence on the extent to which different tasks share a common cognitive process(es) is lacking.

Recently, fMRI studies using a multi-voxel pattern analysis (MVPA) and representational similarity analysis (RSA) approach have compared patterns of activation for self-reference processing to a few other tasks (Yankouskaya et al., 2017; Feng et al., 2018; Parelman et al., 2022). For example, a classifier trained to dissociate positive versus negative affect (Chavez et al., 2017; Yankouskaya et al., 2017) could also distinguish between thinking about self and other in ventral mPFC (vmPFC), suggesting that self-reference processing evokes positive affect. When comparing self-reference to other-reference, MVPA studies have demonstrated distinct patterns of activation in mPFC (Koski et al., 2020; Parelman et al., 2022). Similarly, RSA studies have indicated different patterns of activation in mPFC for thinking about self, close and distant others (Feng et al., 2018; Courtney and Meyer, 2020), as well as for different dimensions of self (Feng et al., 2018). Although these results begin to clarify scholarly understanding of affective and cognitive processes during self-thinking, the degree to which other internally focused processes (namely, introspection and memory) explain self-reference, remains unknown.

Using MVPA, we aim to test similarities and differences between the self-reference task and three other tasks that also rely on internal representation and are known to robustly activate the mPFC. These are the other-reference (Denny et al., 2012; Murray et al., 2012), autobiographical memory (Addis et al., 2007; Summerfield et al., 2009) and introspection tasks (Gusnard et al., 2001; Ochsner et al., 2004; Goldberg et al., 2006; Araujo et al., 2015). Furthermore, using variance partitioning analysis, we aim to quantify how much of explainable variance in mPFC activation patterns during self-thinking can be explained by activation patterns during the other three tasks.

## Method

### Preregistration

We preregistered the sample size, hypotheses, participant exclusion criteria, and data analysis plan at the Open Science Framework (https://osf.io/mn9fz). Unless otherwise noted, we analyzed the data in accord with the preregistration.

### Participants

The experiment was approved by the Kochi University of Technology ethics committee. Before the online autobiographical memory session, participants ticked a box to indicate their consent. We obtained written consent prior to the fMRI experiment.

As preregistered, the final sample comprised 35 Kochi University of Technology students (8 women, 27 men), ranging in age from 18 to 22 years (*M* =19.47, *SD* =1.08). We remunerated them with 2,500 Japanese yen. Participants were right-handed, had no history of psychiatric disorders, and had normal or corrected-to-normal vision. We excluded data from one additional participant due to excessive head movement (preregistered exclusion criteria of >3 mm).

### Experimental procedure

The experiment consisted of two parts: (a) online autobiographical memory survey, and (b) fMRI experiment. The two sessions took place on separate days, 6.97 days apart on average (*SD* = 2.54).

#### Online autobiographical memory session

We adapted the autobiographical memory task from (Wen et al., 2020). Prior to the fMRI scan, we instructed participants to write down 15 autobiographical memories, corresponding to one of 15 events each. These memories should pertain to an event bound to a specific time and context that occurred more than one year ago, but after the age of 10 years. The memories ought to be clear so that participants be in a position to remember the relevant people, objects, and location in detail.

#### Stimuli preparation

We selected for each participant 10 of the 15 listed event memories. We used the selected memories as stimuli in the autobiographical memory task during the fMRI experiment. We based memory selection on the amount of detail and number of characters included in each description. We matched the number of characters with stimuli in the general knowledge task (see below). For each memory, we removed critical words and replaced them with blank underscores prior to the fMRI experiment.

#### The fMRI experiment

The fMRI experiment consisted of the following three tasks (Figure 1): (a) self/other trait judgement task, (b) introspection task, and (c) autobiographical memory task. The self/other trait judgment task had three conditions (Figure 1a-c), whereas the introspection (Figure 1d & 1e) and autobiographical memory (Figure 1f & 1g) tasks had two conditions each. Thus, there was a total of seven conditions. Participants completed five fMRI runs, with each run lasting approximately 6.5 minutes. Each run included two blocks of seven conditions for a total of 14 blocks. We pseudorandomized the block order within each run, so that the same task block was not presented twice in a row. At the beginning of each block, participants viewed a cue for 1 second indicating that the task that was about to commence. All text stimuli were in Japanese. We programmed all tasks in Psychtoolbox (http://psychtoolbox.org/) with Matlab software (version 2018a; http://www.mathworks.co.uk).

**Figure 1.**
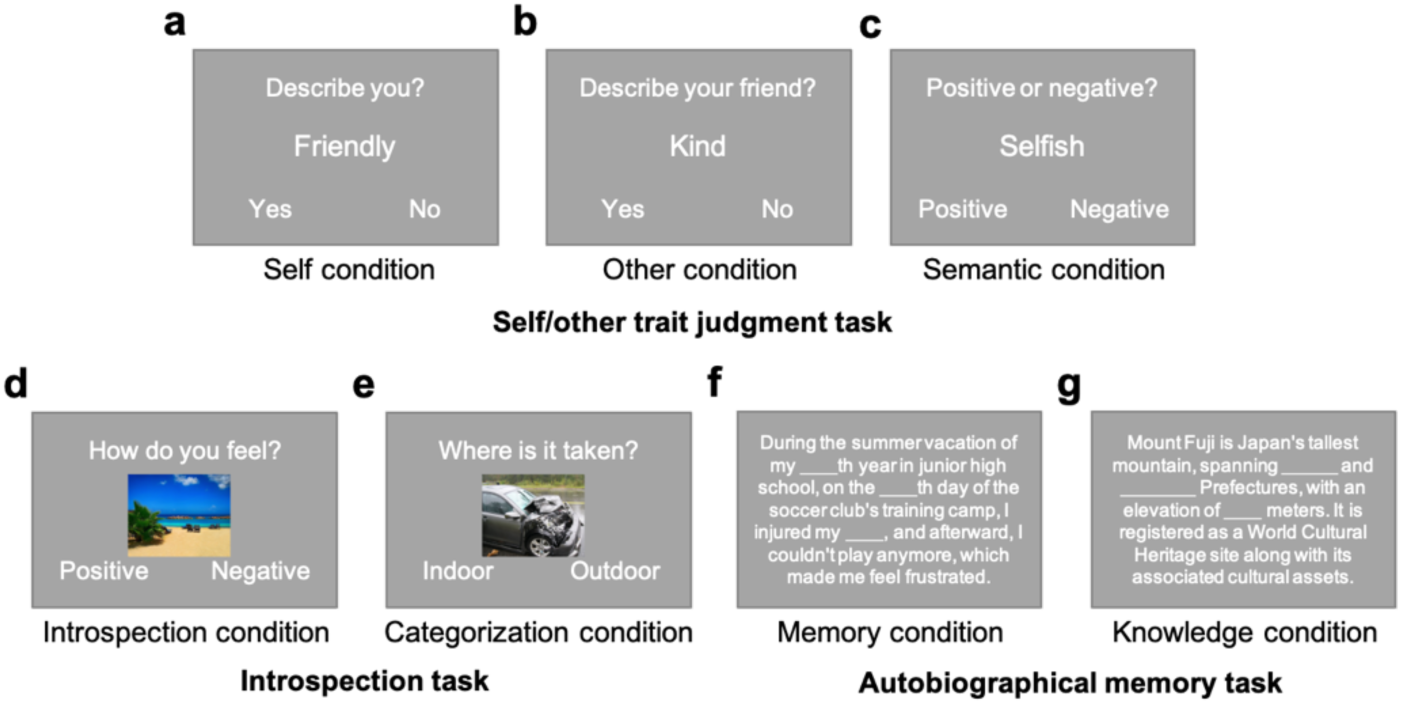
Examples of a trial/block for each of the seven conditions across the three tasks. The self/other trait judgment task consisted of (**a**) self-reference condition, (**b**) other-reference condition, and (**c**) semantic condition. The introspection task consisted of (**d**) introspection task and (**e**) categorization task. The autobiographical memory task consisted of (**f**) memory condition and (**g**) general knowledge condition.

##### Self/other trait judgement task

The stimuli comprised 40 trait adjectives from a pool of normalized trait adjectives (Anderson, 1968), which we translated into Japanese. The stimuli consisted of an equal number of positive (e.g., “honest,” “trustworthy”) and negative (e.g., “mean,” “greedy”) traits. For each trial, we presented a trait in the middle of the screen. In the self-reference block (Figure 1a), we asked participants to judge whether each trait describes them. In the other-reference block (Figure 1b), before fMRI scanning, we asked participants to write down the name of one of their close friends on a piece of paper. During scanning, we instructed them to judge whether each trait describes this specific friend. In the semantic judgment block (Figure 1c), we instructed them to judge whether each trait is positive or negative. We used the same 40 adjectives across the three tasks. We presented each trial for 2 sec, followed by a 1 second fixation cross, and we presented four traits in each block. We randomly determined for each participant the order of traits in each of the self-reference, other-reference, and semantic conditions, but each block always included two positive and two negative words. We presented a fixation cross for 12 sec before the next block.

##### Introspection task

We adapted the introspection task from (Gusnard et al., 2001). It consisted of two conditions: introspection and categorization. We downloaded 40 picture stimuli (i.e., images of objects, animals, or sceneries) from the Open Affective Standardized Image Set (Kurdi et al., 2017). Half of the stimuli were negative and half positive. For each trial in the introspection block (Figure 1d), we presented participants with an image and asked them how the image made them feel. They could respond “positive” or “negative.” In the categorization block (Figure 1e), we asked participants to judge whether each picture depicted a scene that was “indoors” or “outdoors.” We used the same 40 images across the two tasks. For each participant, we randomly determined the order of images in each of the introspection and categorization tasks, but each block always included two positive and two negative images. Each trial lasted for 2 sec, and we displayed a fixation cross for 1 second before the next image appeared. After an introspection block, we displayed a fixation cross for 12 sec before the next block.

##### Autobiographical memory task

The autobiographical memory task (Wen et al., 2020) comprised two conditions: memory and knowledge. For each trial in the memory condition (Figure 1f), participants encountered one of the memories they had previously listed in the online autobiographical memory session. Each memory consisted of, on average, 67.6 Japanese characters (*SD* = 7.25), which we matched with the length of the stimuli used in the knowledge condition. Within each memory, we replaced three critical words with blank underscores. We asked participants to recall the memory and fill in the blanks for the missing words, but do so in their mind rather than by pressing a button (i.e., we recorded no responses during this task).

In the knowledge condition (Figure 1g), we presented participants with text related to general knowledge (*M* = 67.8 characters, *SD* = 8.11 characters), such as a description of a common topic (e.g., Mt. Fuji, football, seatbelt), in which we replaced certain words with blank underscores. We instructed participants to think of appropriate words to fill in the blanks.

In both conditions, we presented each text stimulus for 14 sec and followed it by a fixation cross (4-6 sec). Next, we asked: “Were you recollecting a specific event?” (1 = *not at all*, 5 = *extremely vividly*). Participants had up to 6 sec to respond. We presented a fixation cross for 10 sec before the next block.

## Behavioral data analysis

We conducted a one-way Analysis of Variance to compare reaction time (RT) and response rates across the self-reference, other-reference, and semantic judgment tasks. Given that the RT data were not normally distributed, we log-transformed them beforehand. We followed up significant effects with a Bonferroni-corrected tests. All reported p values were two-sided.

## fMRI data acquisition

We acquired images using a 3.0 T Prisma Siemens MRI scanner with a 64-channel phased-array head coil. For functional imaging, we used T2*-weighted gradient-echo echo-planar imaging (EPI) sequences. We acquired 42 contiguous transaxial slices (covering almost the entire cerebrum) with a thickness of 3 mm, in an interleaved order. We acquired the images with the following parameters: Time repetition (TR) = 2500 ms, echo time (TE) = 25 ms, flip angle (FA) = 90°, field of view (FOV) = 192 mm^2^, matrix = 64 × 64. Additionally, we acquired a T1-weighted structural image (with 1mm isotropic resolution) from each participant.

## fMRI data preprocessing

We carried out preprocessing and statistical analysis in SPM 12 (Welcome Department of Imaging Neuroscience), implemented in MATLAB (Math Works). To allow for T1 equilibration, we discarded the first four volumes before preprocessing and data analyses. We used SPM 12’s preproc_fmri.m script to perform preprocessing of the fMRI data. We spatially realigned all functional images within each run to the mean using 7th-degree B-spline interpolation. We normalized the volumes to MNI space using a transformation matrix that we obtained from the EPI normalization of the first participant to the EPI template. We resampled the volumes to a voxel size of 3 × 3 × 3 mm^3^, that is, we retained the original voxel size. We used the 7th-degree B-spline interpolation option for normalization. We applied spatial smoothing (of 8 mm FWHM) to the data for the whole brain univariate analysis. To maintain fine-grained activation patterns, we did not apply smoothing to the data for representational similarity analysis nor for multivariate pattern analysis.

## Univariate fMRI analysis

### General Linear Model (GLM)

We first ran a conventional GLM analysis, modeling separately each of the seven conditions. We also included six head motion parameters as nuisance regressors. To examine mPFC activation, we created the following six contrast images for each participant: (a) self > semantic, (b) other > semantic, (c) self > other, (d) introspection > categorization, (5) memory > knowledge, (6) rest > semantic + knowledge + categorization. We used the last contrast to identify regions that showed increased activations during passive rest compared to externally focused tasks (Shulman et al., 1997; Gusnard et al., 2001; Wen et al., 2020)). Furthermore, we created the seven additional contrast images (each of the seven tasks relative to the implicit baseline [i.e., rest]). We used the spmT images from these contrasts in the subsequent MVPA and RSA analyses (details below).

### Group analysis

We conducted a second-level whole brain group analysis for each of the contrasts. We set the statistical threshold at *p* < 0.001 voxel-wise (uncorrected) and cluster *p* < 0.05 (FWE corrected for multiple comparisons).

## Representational similarity analysis

We conducted the RSA to test the similarity in activation patterns between the self and each of the other-reference, introspection, and memory conditions. For each participant, we extracted neural data from the spmT image of each contrast, and we computed neural representational similarity matrix (RSM; Figure 2a) based on Pearson correlation across activation patterns in each pair of conditions across the five runs. There are three model RSMs (Figure 2b-d), each of which addresses the similarity between the self and (a) other, (b) introspection, (c) memory. Given that we are interested in the similarity between the self and other, independently of similarities across the remaining conditions, we excluded from analyses the irrelevant conditions. For example, when testing the self = introspection model (Figure 2c), we excluded the other-reference, memory, and knowledge conditions so that pattern similarities involving those irrelevant conditions would not affect the results. We evaluated the fit between the neural RSM and model RSM via Kendall’s tau-a for each participant (Nili et al., 2014).

**Figure 2.**
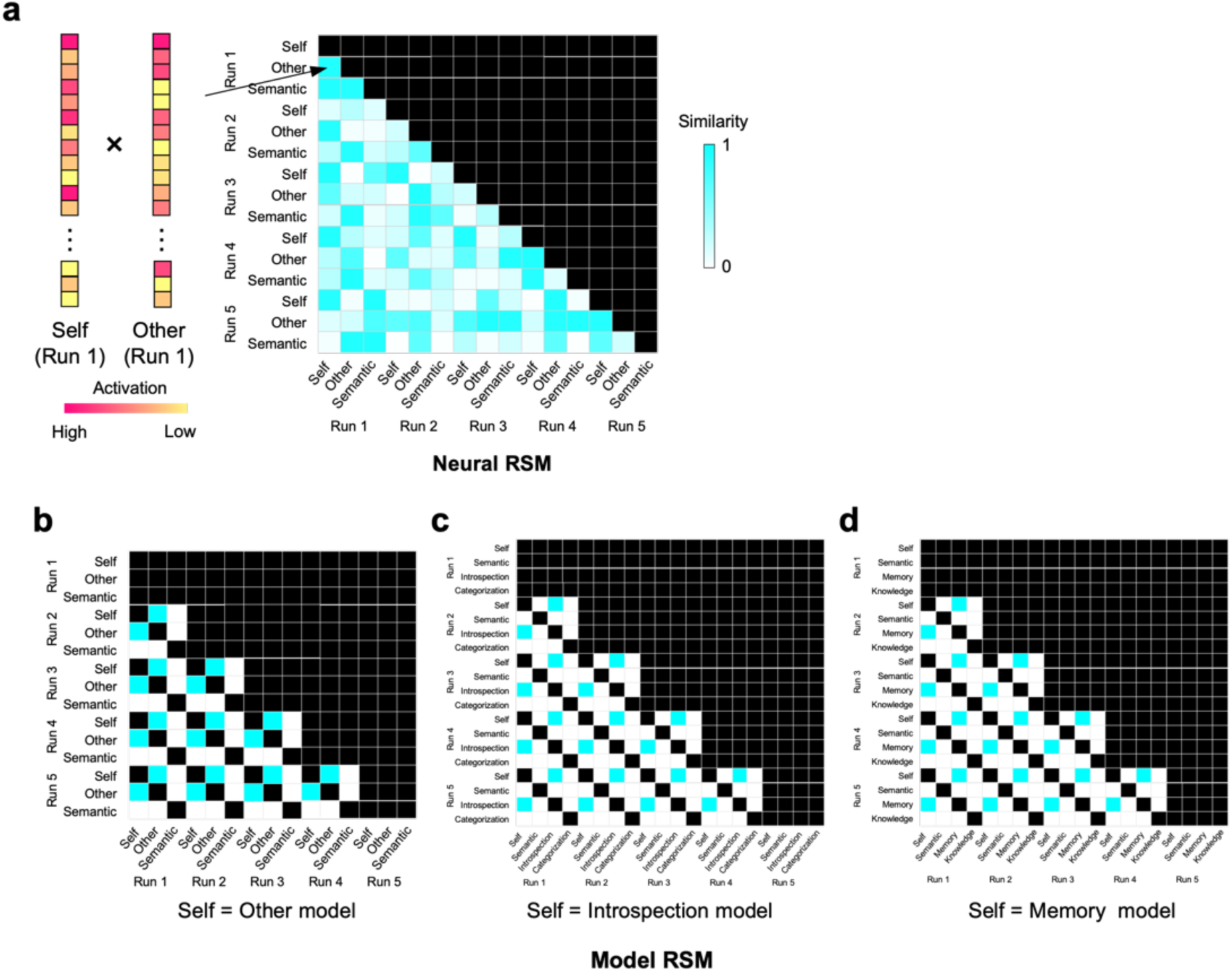
Schematic illustrations of representational similarity analysis (RSA). (**a**) For each participant, we created a neural representational similarity matrix (RSM) by computing Pearson correlations between activation patterns during two tasks across five runs. (**b**) Self = Other model RSM. (**c**) Self = Introspection model RSM. (**d**) self = memory model RSM. In each neural/model RSM, we excluded cells in black from the analysis. In panels **b-d**, cells in cyan represent 1 (similar), whereas cells in white represent 0 (dissimilar). We evaluated fit between the neural versus each model RSMs through Kendall’s tau-a (Nili et al., 2014).

Activations of any two conditions within the same run are likely to be positively correlated largely due to shared physiological noises (e.g., Alink et al., 2015); as such, we excluded correlations between any pairs of conditions within the same run to the model RSM. We also excluded correlations between neural responses of the same conditions (Ritchie et al., 2017). We ran these RSAs using a searchlight approach (explained below).

### Classifier-based MVPA

The above RSA tests whether activation patterns are similar between two conditions. We proceeded to conduct classifier-based MVPA to examine whether activations patterns in the two conditions were distinct. We implemented a linear support vector machine (SVM), carried out via MATLAB in combination with LIBSVM (http://www.csie.ntu.edu.tw/~cjlin/libsvm/) (Wake and Izuma, 2017; Levorsen et al., 2021), with a cost parameter of c = 1 (default).

We used MVPA to find out if the activation patterns for the following contrasts were distinct: (a) self > semantic versus other > semantic, (b) self > semantic versus introspection > categorization, and (c) self > semantic versus memory > knowledge. For each participant, we extracted neural data from the spmT image of each of these contrasts. To evaluate classification performance, we employed a leave-one-run-out cross-validation procedure. Thus, we first left out one run in each cross-validation, and, using the data from the rest of runs, we trained a classifier that discriminates (e.g., activation patterns between self > semantic versus introspection > categorization contrasts). Subsequently, we tested the classifier performance using the data from the left-out run. We repeated this procedure five times leaving out a different run each time, and we averaged the five classification accuracy values. Like the RSA, we ran the classifier-based MVPA using a searchlight approach (below).

### Searchlight analysis

We conducted the RSA and MVPA with a searchlight approach (Kriegeskorte et al., 2006). For the RSA, we extracted local patterns of neural activity from searchlights with a three-voxel radius, so that each searchlight consisted of a maximum of 123 voxels (and less on the edges of the brain). We made a neural RSM from each searchlight and computed Kendall’s tau-a between neural versus each of the three model RSMs (Figure 2), which we saved for a center voxel, resulting in three correlation maps for each participant.

Similarly, for the classifier-based MVPA, we carried out MVPA within each searchlight, and we saved a classification accuracy for a center voxel, resulting in a total of three classification accuracy maps for each participant. Within each searchlight, we removed mean activity by subtracting the mean value of a searchlight sphere from values of the individual voxel so that mean activation difference across conditions could not account for MVPA results.

#### Group analysis

We applied smoothing before the group analysis of the RSA and MVPA outputs (with a Gaussian kernel of 4-mm FWHM). Following the smoothing, we entered the Kendall’s tau-a maps and classification accuracy maps into a second-level permutation-based analysis (with 5,000 permutations). We used the Statistical Non-Parametric Mapping toolbox for SPM (Nichols and Holmes, 2002). Within the preregistered mPFC region of interest (ROI), we set a statistical threshold (i.e., voxel-level) at *p* < 0.005, and a cluster-level threshold at *p* < 0.05 (FWE corrected). Outside of the mPFC, we set a statistical threshold at *p* < 0.001, and a cluster-level threshold at *p* < 0.05 (FWE corrected).

## ROI analysis

We further investigated the role of the mPFC in thinking about the self by running a ROI analysis. We used Neurosynth (https://neurosynth.org/; Yarkoni et al., 2011) to define our mPFC ROI independently of our data. We downloaded an association map (thresholded at q < .01, False Discovery Rate corrected), which we generated from a term-based meta-analysis with the label “self-referential” (downloaded on October 10th, 2023). The mPFC ROI included 308 voxels. We ran the following multivariate pattern regression analyses within the ROI.

## Multivariate pattern regression

The above RSA and classifier-based MVPA address neural pattern similarity and difference separately for each pair of tasks. We conducted a multivariate pattern regression analysis to compare pattern similarity across multiple tasks within the same framework. We ran a multiple regression analysis where activation patterns of the self > semantic contrast were a dependent variable, whereas those of (a) the other > semantic, (b) introspection > categorization, and (c) memory > knowledge contrasts were independent variables (Figure 3).

**Figure 3.**
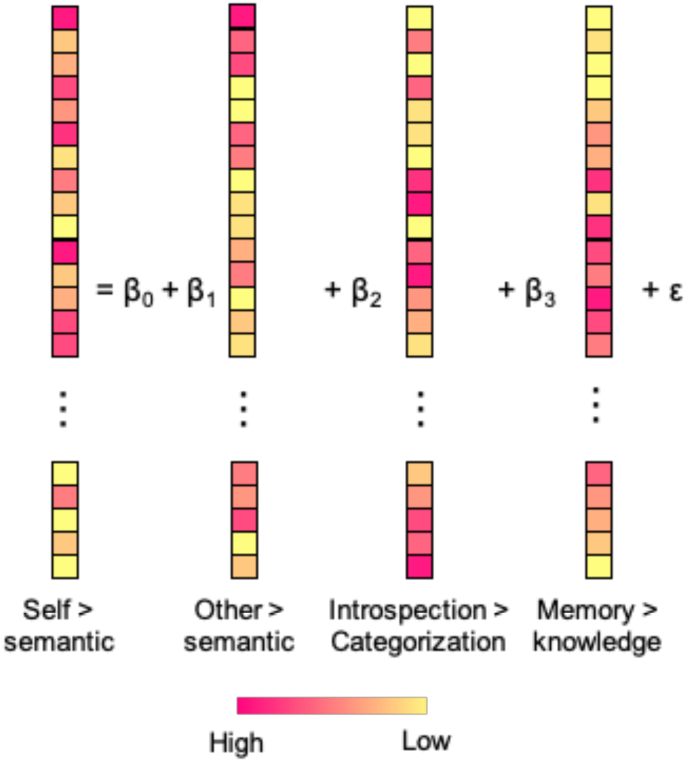
Multivariate pattern regression. Activation patterns of the self > semantic contrast were a dependent variable, whereas activation patterns of the other three contrast were independent variables. Independent and dependent variables were always from different runs.

As stated above, given that activation patterns of any two conditions within the same run are likely to be positively correlated likely due to shared physiological noise, we ran the regression analysis 20 times (all possible pairs of five runs excluding pairs from the same run) so that independent and dependent variables were always from two different runs. We averaged all outputs (beta values and adjusted R^2^) across the 20 regression analyses within each participant.

### Noise ceiling model

To provide an estimate of how much systematic variation in activation patterns of the self > semantic contrast could be explained in the data given measurement noise, we included a noise-ceiling model. This model simply included the data from the self > semantic contrast as both a dependent and independent variable (although they were from different runs) in the multivariate pattern regression. Thus, the only difference between the noise ceiling model and original full model (illustrated in Figure 3) was the inclusion of activation patterns of the self > semantic contrast as another independent variable in the noise ceiling model.

### Variance Partitioning Analysis

Following the multivariate pattern regression analysis, we carried out variance partitioning analysis to infer the amount of unique and shared variance between three different predictors. We conducted seven multiple regression analyses: one with all three independent variables as predictors (illustrated in Figure 3), three with different pairs of two independent variables as predictors, and three with individual independent variables as predictors. Comparing the explained variance (R^2^) of a model used alone with the explained variance when used with other models would allow us to infer the amount of unique and shared variance between different predictors.

### Permutation test

To assess the significance of the findings from the multivariate pattern regression analyses and variance partitioning analysis, we ran permutation tests where voxels were randomly shuffled. The self > semantic contrast and other > semantic contrast have the semantic condition as a common control condition, and this common control condition is likely to bias a beta value associated with the other > semantic activation patterns to a positive direction. Thus, our permutation test randomly shuffled beta activation map of the self-reference condition (i.e., self > implicit rest contrast). We computed a randomly-shuffled-self > semantic contrast image (and a corresponding t-statistics map) so that the effect of the similarity in neural responses between the semantic task versus each of the remaining five tasks remained intact in each permutation. We repeated this step 1,000 times to estimate null distributions. Furthermore, shuffling voxels may overly destroy spatial autocorrelation in the original data, which might bias results of the permutation test. Thus, we smoothed shuffled data via a Gaussian kernel with the standard deviation of 0.86 before conducting a multiple regression analysis (see Burt et al., 2020 for a similar approach). We selected a standard deviation of 0.86, because it produced the smallest sum of square error between the smoothness (quantified as Moran’s I based on an inverse Euclidean distance matrix; Moran, [1950]) of the original data versus that of shuffled-and-then-smoothed data (repeated 1,000 times; we tried all standard deviation values ranging from 0 to 2.0 with an increment of 0.2).

## Deviations from preregistration

We deviated from the preregistration as follows. First, we preregistered and conducted MVPA testing pattern generalizability (i.e., cross-task classification) which, like the RSA, aims to examine the similarity in activation patterns between two conditions. However, we do not report relevant results, because they were similar to the results of the RSA reported below; also, this analysis is inappropriate when testing the similarity between the self-versus other-reference conditions due to their common control condition. Second, we did not preregister the following: behavioral data analyses, reaction time (RT)-controlled MVPA, multivariate pattern regression, and variance partitioning analyses.

## Results

### Behavioral results

During the self/other trait judgment conditions, participants pressed one of the two keys in almost all trials in the self (99.6%), other (99.9%), and semantic (99.9%) conditions. There was a significant difference in RT across the three conditions (F_(2,68)_ = 18.31, *p* < 0.001). Pairwise t-tests revealed that RTs were significantly different from each other across conditions. RTs in the self-reference condition (*M* = 1.21 sec, *SD* = 0.18 sec) were significantly longer than those in the other-reference condition (*M* = 1.14 sec, *SD* = 0.25 sec; *p_corrected_* = 0.004) and in the semantic condition (*M* = 1.08 sec, *SD* = 0.19 sec; *p_corrected_* < 0.001). RTs in the other condition were significantly longer than those in the semantic condition (*p_corrected_* = 0.026).

We next examined if RTs in the other-reference condition were influenced by response similarity between the self and other, as reported in a previous study (Thornton and Mitchell, 2018). We ran a multiple regression analysis with RT in the other-reference condition as a dependent variable, and response similarity as an independent variable (1 = same responses to the same trait, -1 = different responses). We also entered as independent variables participant response (1 = yes, -1 = no), trait valence (1 = positive, -1 = negative), number of characters of each word stimulus, and the interaction between participant response and trait valence. We obtained a significant effect of response similarity (*t*_(34)_ = 3.79, *p* = 0.003). RTs were shorter when the self- and other-reference judgments for the same trait were identical (i.e., both yes or both no). Although this result suggests egocentric anchoring and adjustment in other-reference judgment, we observed a similar effect in the self-reference condition (see below). Number of characters was significantly related to RTs, meaning the more characters a word had, the slower the participant responded (*t*_(34)_ = 3.80, *p* = 0.003). We also obtained a significant Participant Response × Trait Valence interaction (*t*_(34)_ = 4.42, *p* = 0.005). Participants were slower to respond yes than no when judging if a negative trait described their friend, whereas they did not differ in their responses to positive traits. No other significant effect emerged.

We conducted the same regression analysis for the self-reference condition to test if RTs in the self-reference condition were influenced by response similarity between the self and other. We found only a significant effect of number of characters (*t*_(34)_ = 2.97, *p* = 0.027). The effect for response similarity was trending (*t*_(34)_ = 2.51, *p* = 0.086). When we compared beta values for the self-versus other-reference conditions, we observed no significant difference (*t*_(34)_ = 1.03, uncorrected *p* = 0.31), suggesting that the significant effect of the response similarity obtained in the other-reference condition might be at least partially explained by unknown stimulus features.

Consistent with prior research (e.g., Moran et al., 2006), participants were more likely to endorse a positive trait as self-descriptive and a negative trait as not self-descriptive (*t*_(34)_ = 4.28, p < 0.001). However, we observed this positivity bias in the other-reference condition as well (*t*_(34)_ = 8.46, *p* < 0.001); indeed, this bias was stronger for the other-reference than the self-reference condition, indicating that participants were more other-enhancing than self-enhancing (*t*_(34)_ = 3.48, *p* = 0.001). These results are largely consistent with some findings suggesting that self-enhancement is weaker for East Asian compared to Western individuals (Heine and Hamamura, 2007; but see also Cai et al., 2016).

During the introspection task, participants pressed one of the two keys in almost all trials in the introspection (99.9%) and categorization (99.7%) conditions. RTs during the introspection condition (*M* = 1.08 sec, *SD* = 0.21) were significantly slower than those during the categorization (*M* = 1.14 sec, *SD* = 0.17) condition (*t*_(34)_ = 3.79, *p* < 0.001), likely because some pictures were ambiguous as to whether they were taken indoors or outdoors.

During the autobiographical memory task, participants successfully gave their vividness rating within the time limit of 6 sec for almost all trials in the memory (99.0%) and knowledge (98.7%) conditions. Vividness ratings were significantly higher in the memory (*M* = 4.37, *SD* = 0.40) compared to the knowledge (*M* = 2.45, *SD* = 0.79) condition (*t*_(34)_ = 17.73, *p* < 0.001), testifying to the effectiveness of our memory manipulation.

## fMRI results

### Univariate analysis results

Replicating findings from several studies (Denny et al., 2012; Murray et al., 2012), the self > semantic contrast significantly activated the midline structure including mPFC and PCC (Figure 4a and Supplementary Table 1). The other > semantic contrast activated similar regions (Figure 4b and Supplementary Table 2). Left temporoparietal junction (TPJ) was also commonly activated by the self and other conditions. Although there were some regions that were uniquely activated by either the self > semantic or other > semantic contrast when the self and other conditions were directly compared, the self > other contrast did not lead to any significant activation. The opposite contrast (other > self) revealed only one significant cluster in PCC (303 voxels; x = 6, y = -64, z = 29).

**Figure 4.**
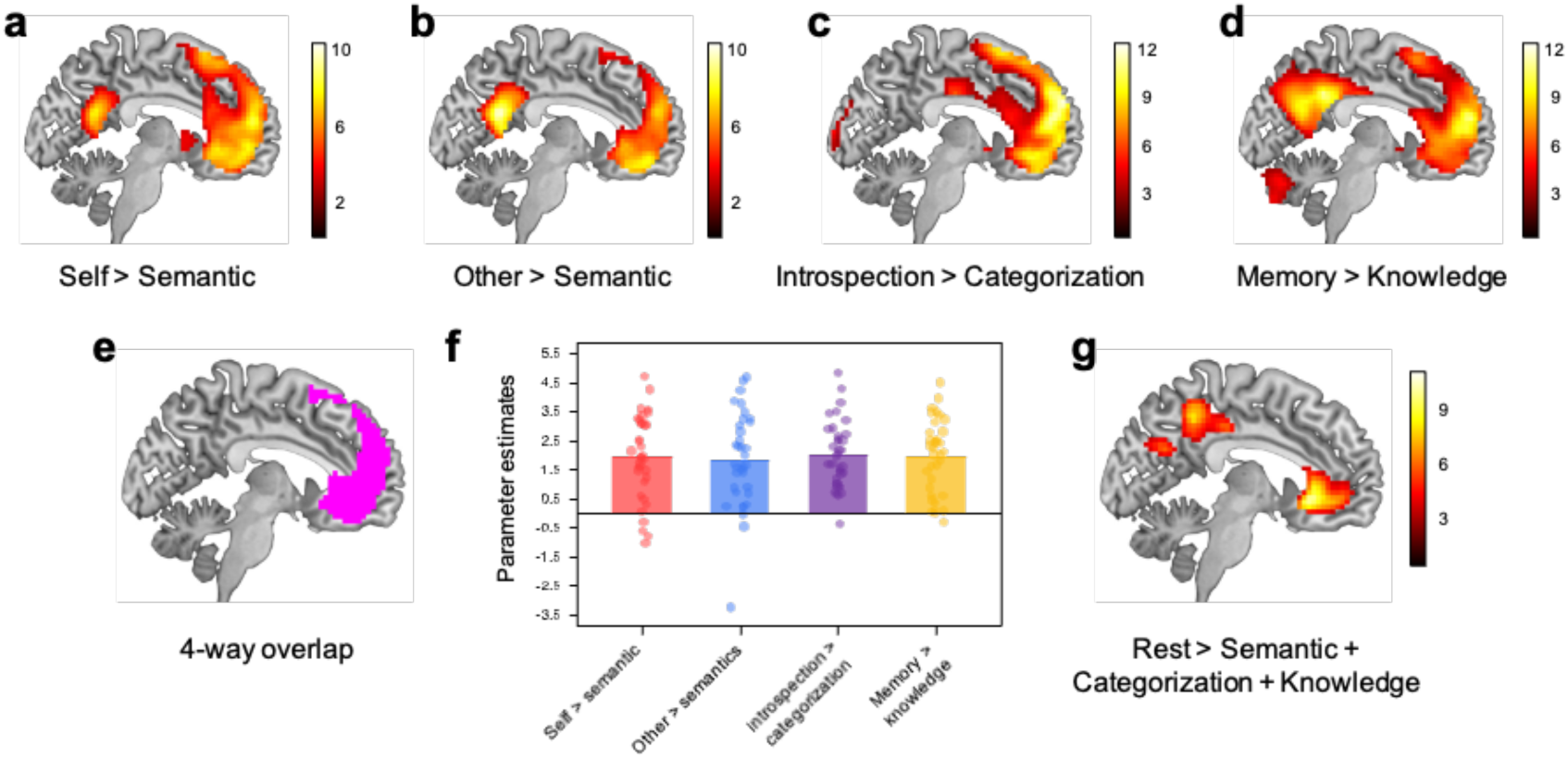
Sagittal slices (x = -6) showing results of the univariate analyses. (**a**) Areas significantly activated by the self > semantic contrast. (**b**) Areas significantly activated by the other > semantic contrast. (**c**) Areas significantly activated by the introspection > categorization contrast. (**d**) Areas significantly activated by the memory > knowledge contrast. (**e**) Areas commonly activated by the all four contrasts (1,565 voxels). Only mPFC showed significant 4-way overlap. For display purposes, we set statistical threshold at *p* < 0.005 and cluster-*p* < 0.05 (FWE corrected). (**f**) Parameter estimates of the four contrasts within the mPFC areas commonly activated by the four contrasts (panel e). (**g**) Areas significantly activated by the rest > semantic + categorization + knowledge contrast.

As per prior studies (Gusnard et al., 2001; Goldberg et al., 2006), the introspection > categorization contrast significantly activated the mPFC (Figure 4c). Other activated areas included anterior cingulate cortex, temporal pole, lateral temporal cortex, and lateral occipital cortex (Supplementary Table 3).

The memory versus knowledge contrast significantly activated regions previously implicated in autographical memory including the mPFC, PCC/precuneus, posterior inferior parietal lobule (pIPL), and lateral temporal cortex (LTC) (Kim, 2012; Martinelli et al., 2013) (Figure 4d and Supplementary Table 4).

Taken together, the above four contrasts all significantly activated the common region within the mPFC (1,565 voxels; Figure 4e). Bilateral temporal poles were also commonly activated by all four contrasts (left x = -36, y = 17, z = -22, 109 voxels; right x = 30, y = 14, z = - 22, 185 voxels). No other region was commonly activated. Yet, although the introspection > categorization contrast activated the PCC (92 voxels), it did not pass our preregistered cluster-level threshold. When we directly compared the four contrasts to each other within the commonly activated mPFC areas, no significant difference emerged (*F*_(2.38, 80.86)_ = 0.08, *p* = 0.94; Figure 4f).

Consistent with previous findings (Shulman et al., 1997; Gusnard et al., 2001; Wen et al., 2020), the rest > semantic + categorization + knowledge contrast revealed that areas in the default mode network, including mPFC, PCC, IPL, TPJ/AG, and LTC, were active during rest compared to the externally focused tasks (Figure 4g).

### Results of RSA: Are activation patterns evoked by two tasks similar?

The RSA (Figure 2) aims to test whether the self-reference judgment evoked similar activation patterns with each of the other-reference judgment, introspection, and autobiographical memory.

The Self = Other model (Figure 2b) was significantly associated with a network of brain regions involved in self-reference and social cognition including mPFC, PCC/precuneus, bilateral inferior frontal gyrus (IFG), bilateral superior temporal sulcus, and bilateral temporal pole (Figure 5a). We report in Supplementary Table 5 all areas significantly associated with the Self = Other model. However, the other two models were associated only with mPFC and left IFG. In particular, the Self = Introspection model (Figure 2c and Supplementary Table 6) was significantly associated with mPFC (x = 0, y = 53, z = 35, 1,930 voxels; see Figure 5b) and left IFG (extending to temporal pole; x = -51, y = 20, z = 2, 379 voxels). Further, the Self = Memory model was significantly associated with mPFC (x = -12, y = 44, z = 5, 103 voxels; Figure 5c and Supplementary Table 7) and left IFG (x = -45, y = 26, z = -7, 368 voxels).

**Figure 5.**
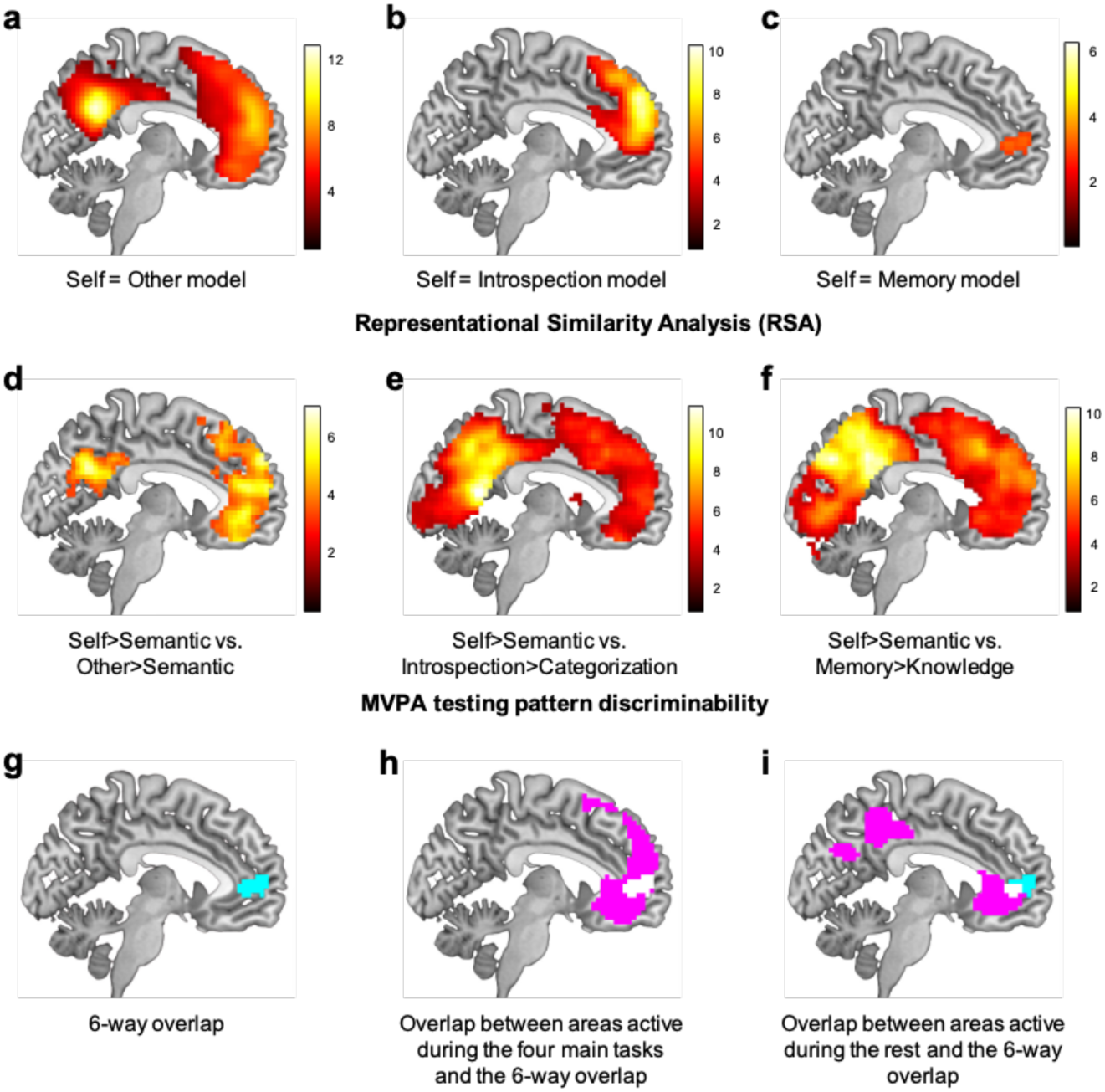
(**a-c**) Sagittal slices (x = -6) showing results from the RSA. Significant areas indicate that activation patterns of the two contrasts were similar. (**d-f**) Sagittal slices (x = -6) showing results from the MVPA testing pattern discriminability. Significant areas indicate that activation patterns of the two contrasts were distinguishable. For display purposes, we set statistical threshold at *p* < 0.005 and cluster-*p* < 0.05 (FWE corrected). (**g**) A sagittal slice (x = -6) showing the mPFC area that showed 6-way overlap (overlap across areas shown in panel a-f). (**h**) A sagittal slice (x = -6) showing overlap between univariate and MVPA results. Magenta represents areas activated commonly by the four univariate contrasts (Figure 4e), and white represents 6-way overlapped region depicted in panel **g**. (**i**) A sagittal slice (x = -6) showing overlap (white areas) between areas activated by the rest > semantic + categorization + knowledge contrast (magenta; Figure 4g) and the 6-way overlapped region depicted in panel **g** (cyan).

### Results of MVPA testing pattern discriminability: Are activation patterns evoked by two tasks distinguishable?

The classifier-based MVPA tested pattern discriminability with a searchlight approach. It addressed whether activation patterns evoked by different tasks were distinguishable or linked to different cognitive processes. Indeed, activation patterns evoked during the self-reference task (relative to the semantic task) were distinguishable from the other-reference task in the mPFC, PCC, and right superior temporal sulcus (extending to the temporal pole; Figure 5d and Supplementary Table 8). These areas largely overlapped with the areas activated by the self- and other-reference condition relative to the semantic condition (Figure 4a and 4b), indicating that those areas were commonly activated both by the self and other conditions compared to the semantic condition, but their activation patterns were systematically different. Given that the self and other conditions had the semantic condition as common control, the difference between the self and other conditions is likely to be underestimated in this analysis.

In contrast, activation patterns elicited by the self-reference condition were distinguishable from each of the introspection and memory conditions in a number of regions across the whole brain including the mPFC, PCC, intraparietal lobule, middle temporal gyrus, and TPJ (Figures 5e and 5f, Supplementary Tables 9 and 10).

These results, together with the RSA results reported above, indicate that mPFC activation patterns during the self-reference judgement were similar to those elicited during each of the other-reference judgement, introspection, and autobiographical memory (Figure 5a-c). Nonetheless, they were still distinguishable from activation patterns of each of the three tasks (Figure 5d-f). In fact, there was one cluster within the mPFC (Figure 5g; a total of 96 voxels) showing significant association/classification accuracy in all six analyses (Figure 5a-f), and the mPFC cluster is the only region that showed the 6-way overlap with the cluster size larger than 20 voxels. This 6-way overlap was located in the anterior part of the mPFC (Brodmann Area [BA] 10) and pregenual anterior cingulate cortex (pgACC; BA 32). Furthermore, this cluster entirely overlapped with the areas commonly activated by the four contrasts in the univariate analyses (Figure 5e). It also showed a substantial overlap (53 out of 96 voxels [55.2%]) with the areas significantly active during the rest (Figure 5g), indicating that most of the 6-way overlap area (Figure 5e) is located in the mPFC within the default mode network.

### Results of ROI analyses

We conducted additional ROI analyses to refine the findings and run control analyses. We defined the mPFC ROI independently of our own data using Neurosynth (see Methods). This ROI analysis focused on areas within the mPFC most strongly associated with self-reference processing (Figure 6a).

**Figure 6.**
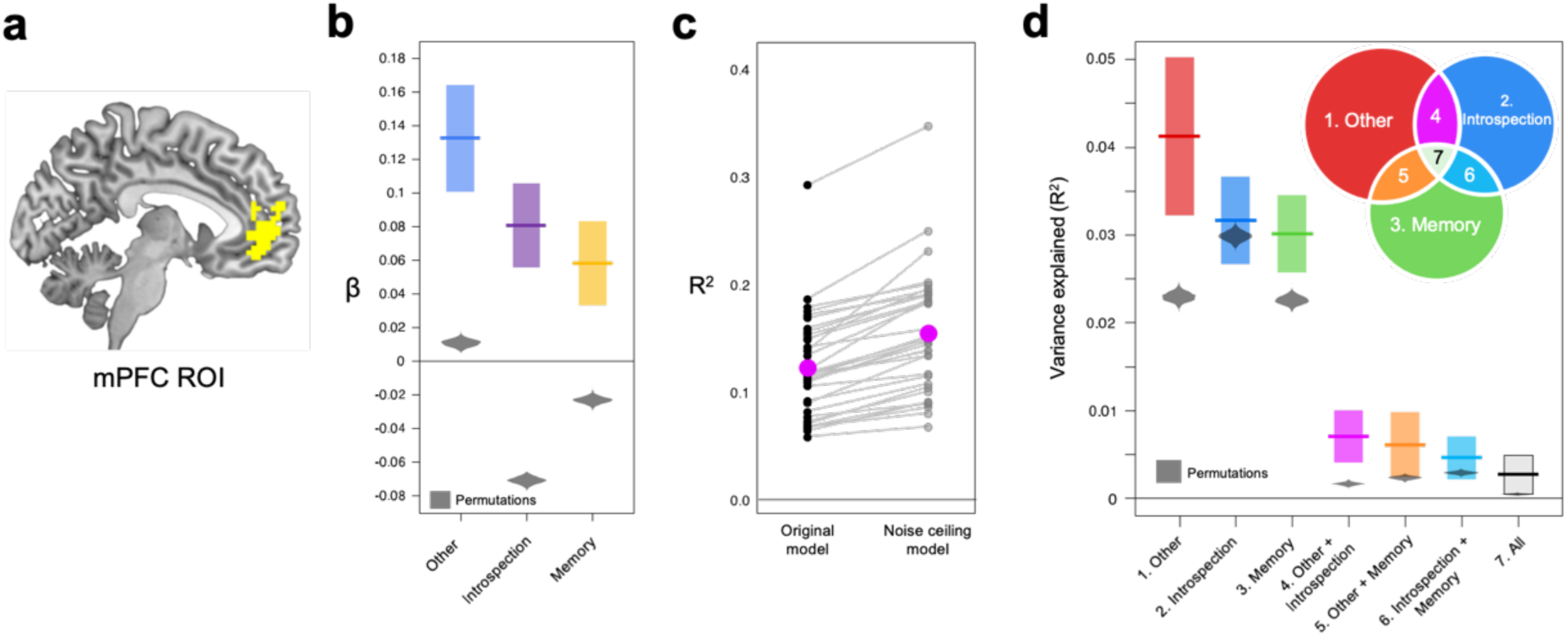
Results of the multivariate pattern regressions. (**a**) A sagittal slice (x = -6) showing the mPFC areas used in the ROI analysis. We defined the mPFC ROI with the term “self-referential” based on Neurosynth term-based meta-analysis. (**b**) Beta values from the multivariate pattern regression with activation patterns of the self > semantic contrast as a dependent variable. Colored horizontal lines indicate mean beta values, and lower/upper box limits represent 95% confidence intervals (CIs). (**c**) Adjusted R^2^ from the original regression model (Figure 3) and the noise ceiling model. Pink circles indicate mean R^2^, and black/grey circles indicate R^2^ of individual subjects. (**d**) Variance in mPFC ROI activation patterns of the self-reference condition that was explained by activation patterns of the other, introspection, and memory conditions. In panels **b** and **d**, bell shaped gray areas indicate permutation distribution.

*Does the difference in activation patterns between the self- and other-reference conditions simply reflect the difference in RTs between them*? According to our behavioral results, RTs were significantly longer for the self-reference condition compared to the other-reference condition. Thus, the difference in activation patterns between the two conditions might be explained by the difference in RTs (e.g., task difficulty). To rule out this possibility, we ran additional GLM where we categorized self- and other-reference task blocks into short and long RT blocks based on average RTs in each block. We modeled the other five tasks in the same way as the original GLM. Then, we ran an MVPA analysis testing whether it can distinguish activation patterns of the mPFC ROI during the short versus long RT blocks.

Within the mPFC ROI, the average accuracy for classifying the short and long RT blocks was 51.71%, which did not differ significantly from the theoretical chance level of 50% (Wilcoxon signed rank test, *p* = 0.31). Also, it was significantly lower than the accuracy for classifying actual self-versus other-reference blocks (average = 63.14%; paired-sample Wilcoxon signed rank test, *p* = 0.002). We additionally ran the same MVPA (short vs. long RT blocks) across the whole brain with a searchlight approach, but did not find any significant area. Taken together, the difference in RTs between the self and other conditions is unlikely to explain the difference in activation patterns between the two conditions.

*Which task best explains activation patterns of the self-reference condition*? The results of the RSA reported above (Figure 5a-c) indicate that mPFC activation patterns during the self-reference condition were similar to those of the other-reference, introspection, and memory conditions. However, these analyses addressed neural pattern similarity separately for each pair of tasks. To compare pattern similarity across three tasks within the same framework, we carried out a multivariate pattern regression analysis where activation patterns of the self > semantic contrast were a dependent variable, whereas those of the other > semantic, introspection > categorization, and memory > knowledge contrasts were independent variables (Figure 3). Activation patterns of each of the three contrasts were significantly associated with mPFC ROI activation patterns of the self > semantic contrast (Figure 6b; all *p_perm_* < 0.001), suggesting that the similarity in mPFC neural responses between the self-reference task and each of the other three tasks remain significant even after controlling for the effect of neural responses during the other two tasks.

Adjusted R^2^ were significantly lower than the that of the noise ceiling model (*p_perm_* < 0.001; Figure 6c). Hence, there was still unexplained variance even after considering noise in the fMRI data, suggesting that there were patterns of activations specific to the self-reference judgment (not shared by the other three tasks). Activation patterns of the other > semantic, introspection > categorization, and memory > knowledge contrasts collectively explained, on average, 79.44 % of explainable variance in the mPFC activation patterns of the self > semantic contrast.

#### Variance Partitioning Analysis (VPA)

We conducted a variance partitioning analysis to quantify how much variance in the mPFC ROI responses of the self > semantic contrast is explained uniquely by activation patterns of each of the other > semantic, introspection > categorization, and memory > knowledge contrasts, while considered together with the other two conditions. We present the results in Figure 6d. Each of the seven portions significantly explained the variance in neural responses of the self > semantic contrast (all *p_perm_* < 0.001). The results suggest that mPFC activation patterns reflect multiple cognitive processes. For example, a significant amount of variance explained by all three contrasts indicates that there were specific patterns of mPFC neural responses shared across the self-reference, other-reference, introspection, and memory tasks, which likely reflects a cognitive process common to all tasks. Similarly, a significant amount of variance explained by the other-reference and introspection conditions indicates that there were specific patterns of mPFC neural responses shared across the self-reference, other-reference, and introspection tasks, which likely reflects a cognitive process common to these tasks, but not to the memory task (see below for more discussion).

We ran the same multivariate pattern regression and variance partitioning analyses in other regions related to self-reference (based on the same Neurosynth meta-analysis map with the term “self-referential”) and to the default mode network (based on Andrews-Hanna et al., 2010). The anterior and dorsal parts of the mPFC (amPFC and dmPFC) of the default mode network were the only regions that evinced the same pattern of the results as the mPFC reported above: (a) significantly positive beta values for all three independent variables (multivariate pattern regression), and (b) significantly positive variance explained for all seven portions (variance partitioning analysis; Supplementary Figure 1), indicating a unique and complex role played by the mPFC during thinking about the self.

## Discussion

We provided a more nuanced and precise picture of the mPFC’s role during thinking about the self. Replicating prior findings, each of the self-reference, other-reference, introspection, and autographical memory tasks activated the mPFC compared to their corresponding control condition (Figure 4). Furthermore, we demonstrated that the relationship between activation patterns during the self-reference task and those of the other three tasks (other-reference, introspection, and autobiographical memory) was intricate. That is, mPFC neural responses during the self-reference task were not simply similar to one task and different from the other two tasks. Instead, the mPFC neural responses during the self-reference task were both similar and distinct at the same time from each of the other-reference, introspection, and autobiographical memory tasks (Figure 5). The mPFC was the only region across the whole brain that evinced these result patterns.

Furthermore, the multivariate pattern regression together with the variance partitioning analyses revealed complex relationships of activation patterns of each of the three other tasks to mPFC neural responses, during the self-reference task (Figure 6). According to the variance partitioning analyses, not only each of the other-reference, introspection, and memory tasks uniquely explained substantial variance in mPFC neural responses during the self-reference task, but also each pair of these tasks and all three tasks jointly explained significant amounts of variance of the mPFC neural responses during the self-reference task (Figure 6d). Hence, it suggests that there are cognitive processes common to thinking about the self and (a) each of the three tasks, (b) each pair of the three tasks, and (c) all three tasks (thus a total of seven different cognitive processes; Table 1). In addition, adjusted R^2^ of the full model (Figure 3) were significantly lower than the that of the noise ceiling model (Figure 6c), suggesting that there are mPFC neural responses (i.e., a cognitive process) specific to the self-reference task. Overall, our results indicate that there are at least eight cognitive processes (i.e., seven cognitive processes listed in Table 1 plus a self-specific process) at play simultaneously when performing the self-reference task, some of which are common across tasks. Our study does not specify what these cognitive processes are (see Table 1 for ideas on possible candidate processes), leaving this issue open for future research. Nonetheless, as to the self-specific cognitive process, in a prior investigation we reported that the self-specific activation patterns depend on the importance of the stimuli for self-concept (Levorsen et al., 2023), and so access to this self-concept information stored in the mPFC may be responsible for the self-specific mPFC activation patterns we presently observed.

**Table 1.**
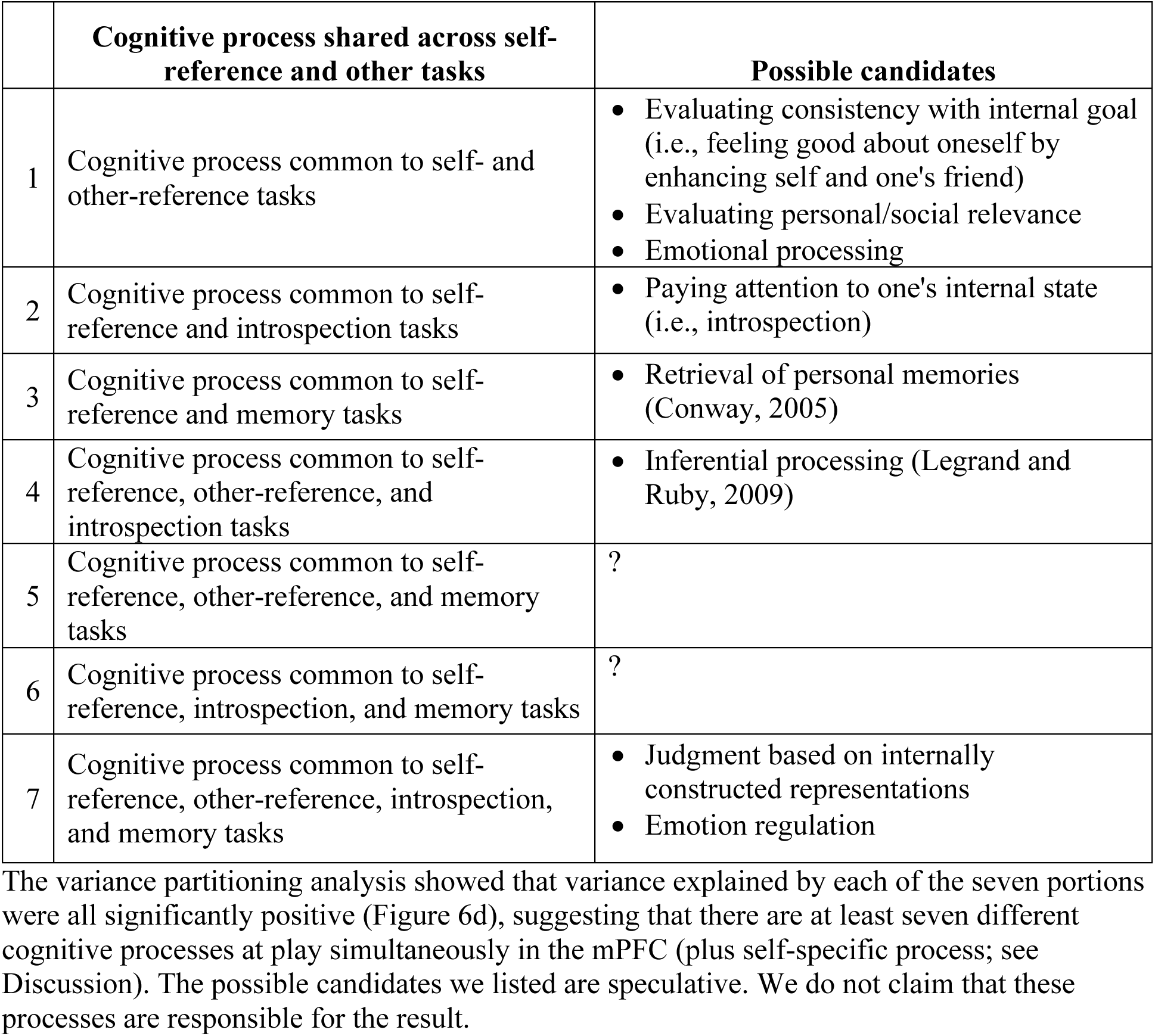
Seven cognitive processes shared across self-reference and other tasks, and their possible candidates

|  | **Cognitive process shared across self-reference and other tasks** | **Possible candidates** |
| --- | --- | --- |
| 1 | Cognitive process common to self- and other-reference tasks | • Evaluating consistency with internal goal (i.e., feeling good about oneself by enhancing self and one's friend)<br>• Evaluating personal/social relevance<br>• Emotional processing |
| 2 | Cognitive process common to self-reference and introspection tasks | • Paying attention to one's internal state (i.e., introspection) |
| 3 | Cognitive process common to self-reference and memory tasks | • Retrieval of personal memories (Conway, 2005) |
| 4 | Cognitive process common to self-reference, other-reference, and introspection tasks | • Inferential processing (Legrand and Ruby, 2009) |
| 5 | Cognitive process common to self-reference, other-reference, and memory tasks | ? |
| 6 | Cognitive process common to self-reference, introspection, and memory tasks | ? |
| 7 | Cognitive process common to self-reference, other-reference, introspection, and memory tasks | • Judgment based on internally constructed representations<br>• Emotion regulation |

This view of the mPFC’s role in the self-reference task invites re-interpretation of prior findings. For example, a few studies showed that mPFC activation patterns are different depending on the target person during the self/other-reference tasks (e.g., self vs. close-other vs. distant other) and dimensions of person knowledge (e.g., traits, physical attributes, social roles) (Feng et al., 2018; Courtney and Meyer, 2020; Koski et al., 2020). The present study suggests that what drives these different mPFC neural responses might be differences in how much each task relies on different information (thus, cognitive processes). For instance, thinking about close others and acquaintances might rely more on one’s autobiographical memory, whereas thinking about unfamiliar others (e.g., celebrities) might rely more on semantic memory (Courtney and Meyer, 2020). The mPFC activation patterns are also likely to vary depending on whether a context is general or specific (I am friendly in general vs. at the university) (Martial et al., 2018) and differences in various dimensions of distance similarity (e.g., temporal, spatial, social, hypothetical) (Tamir and Mitchell, 2011) as these judgements likely rely on distinct sources of information.

Moreover, our variance partitioning analysis demonstrated that, among regions related to self-reference and regions in the default mode network, the mPFC is the only region that had significantly positive variance explained for all seven portions (Figure 6d and Supplementary Figure 1). This result suggests that the mPFC, one of the core hubs of the default mode network (Andrews-Hanna et al., 2010; Andrews-Hanna, 2012), might be a place where necessary information is gathered and integrated for judgments based on internally constructed representations. As a metaphor, to make a soup, one needs to gather ingredients from different parts of the kitchen and mix them in a pot, with different soups often having some common ingredients (e.g., Italian minestrone and Japanese miso soup commonly use some vegetables, and all soups use water). Similarly, to perform a task that requires a decision based on internally constructed representations (Nakao et al., 2012; Andrews-Hanna et al., 2014; Buckner and DiNicola, 2019; Wen et al., 2020; Menon, 2023), necessary information is gathered from different parts of the brain and integrated in the mPFC. Just like Italian minestrone and Japanese miso soup, different tasks often rely on common cognitive processes (e.g., autobiographical memory and introspection), and there may be a common cognitive process(es) for all such tasks, like water for all soups. This view offers an insight into why diverse social and cognitive tasks activate the mPFC. The mPFC has been consistently implicated not only in the four tasks (self-reference, other-reference, introspection, and autobiographical memory) used in the present study, but also in other tasks such as theory of mind, episodic future thinking, and spatial navigation (Andrews-Hanna et al., 2014; Buckner and DiNicola, 2019; Wen et al., 2020; Menon, 2023). It is likely that to perform these tasks one needs to gather information from different part of the brain so as to construct internal representations.

This idea for the role of the mPFC role is largely consistent with roles of the default mode network proposed previously. For example, Yeshurun et al. (2021) considered the default mode network as an active and dynamic sense-making network that integrates incoming extrinsic information with prior intrinsic information to form rich, context-dependent models of situations as they unfold over time. More recently, Menon (2023) argued that the default mode network integrates multiple cognitive functions to create a coherent internal narrative of our experiences. Within these large frameworks on the default mode network function (see also Koban et al., 2021), the present study provides evidence that, among regions in the default mode network, the mPFC is a place where all information converges and is integrated to form coherent internal representation for making a task-relevant judgement. Put otherwise, multiple cognitive processes are performed in the mPFC during a single task (i.e., self-reference task).

Our results highlight an important conceptual challenge for social neuroscientists: Each of many tasks used in social neuroscience involves multiple cognitive processes (or operations), and each of these processes needs to be identified to fully understand the function of the mPFC (and other brain regions) (note that this point is well recognized by memory researchers [e.g., Moscovitch, 1992]). For example, our findings indicate that the difference between the self-reference and semantic tasks pertains not only to the level of self-referential processing, but also to the fact that there are several other additional cognitive processes involved in the self-reference task, some of which are shared with other-reference, introspection, and memory tasks (Table 1). Thus, instead of a traditional brain mapping approach (illustrating which regions are activated by the self-reference > semantic contrast), brain mapping is needed at a much finer scale, linking basic cognitive processes not to univariate activation, but to specific activation *patterns* within a region. Although the utility of a multivariate approach over a univariate approach has been well recognized (and its methodology has been well developed) (Haynes and Rees, 2006; Haxby, 2012), identifying each of various basic cognitive processes involved in a social/cognitive task remains a challenge (e.g., see Schaafsma et al., 2015). The multivariate pattern regression approach (with the variance partitioning analysis), where both dependent and independent variables are neural responses (Figure 3), is a promising approach, as it helps us to statistically decompose complex social/cognitive tasks and identify whether there are unique or common processes across different tasks.

In conclusion, the current findings enhance understanding of the mPFC and its involvement in self-referential thinking by demonstrating its unique role in integrating diverse cognitive processes. The mPFC is not merely activated by self-reference, but also shows complex activation patterns that are both similar and distinct from other cognitive tasks such as other-reference, introspection, and autobiographical memory. Taken together with the role of the mPFC within the default mode network reported previously, the findings indicate that the mPFC serves as a hub where information from various brain regions is gathered and integrated, facilitating tasks that involve constructing internal representations.

## Acknowledgements

This research was supported by a Japan Society for Promotion of Science (JSPS) KAKENHI Grant Number JP19K24680 (to K.I.).

## Notes

### Competing Interest Statement

The authors have declared no competing interest.

