## Supplemental Materials for "Decomposing Cognitive Processes in the mPFC During Self-Thinking"

| | region | MNI coordinate | | | voxels | Multivariate Regression | | | Variance Partitioning Analysis | | | | | | |
|---|---|---|---|---|---|---|---|---|---|---|---|---|---|---|---|
| | | x | y | z | | Other-ref | Introspection | Memory | 1. Other | 2. Introspection | 3. Memory | 4. Other + Introspection | 5. Other + memory | 6. Introspection + memory | 7. All |
| Self-related regions (Neurosynth) | dmPFC | -6 | 41 | 41 | 56 | *** | *** | *** | *** | *** | *** | *** | *** | *n.s.* | *** |
| | PCC | -3 | -58 | 26 | 131 | *** | *** | ** | *** | *n.s.* | *** | *** | *** | *** | *** |
| | Left TPJ | -48 | -64 | 29 | 66 | *** | *** | *n.s.* | *** | *n.s.* | *n.s.* | *** | *** | *n.s.* | *** |
| | Left TP | -60 | -13 | -22 | 70 | *** | *** | *n.s.* | *** | *** | *** | *** | *** | *** | *n.s.* |
| Default mode network (Andrews-Hanna et al., 2010) | vmPFC | 0 | 26 | -18 | 39 | *** | *** | *n.s.* | * | ** | *n.s.* | *n.s.* | * | *** | *n.s.* |
| | amPFC | -6 | 52 | -2 | 93 | *** | *** | *** | *** | ** | *** | *** | *** | ** | *** |
| | dmPFC | 0 | 52 | 26 | 116 | *** | *** | *** | *** | *** | *** | *** | *** | *** | *** |
| | Left TC | -60 | -24 | -18 | 117 | *** | *** | *n.s.* | *** | ** | *** | *** | *** | *** | *n.s.* |
| | Left TPJ | -54 | -54 | 28 | 110 | *** | *** | *n.s.* | *** | *n.s.* | *n.s.* | *** | *** | *n.s.* | *** |
| | PCC | -8 | -56 | 26 | 116 | *** | *** | ** | *** | *n.s.* | *** | *** | *** | *** | *** |
| | PHC | -28 | -40 | -12 | 79 | *n.s.* | *** | *n.s.* | *n.s.* | *n.s.* | *n.s.* | *n.s.* | *n.s.* | *n.s.* | *n.s.* |
| | pIPL | -44 | -74 | 32 | 78 | *** | *** | *n.s.* | *** | *n.s.* | *n.s.* | *** | *** | *n.s.* | *** |
| | Rsp | -14 | -52 | 8 | 83 | *n.s.* | *** | *n.s.* | *n.s.* | *n.s.* | *n.s.* | *n.s.* | *n.s.* | *n.s.* | *n.s.* |
| | TempP | -50 | 14 | -40 | 21 | *n.s.* | *** | *** | * | ** | *n.s.* | *n.s.* | *n.s.* | *n.s.* | *n.s.* |

**Supplementary Figure 1.** Results of the multivariate pattern regressions and variance partitioning analyses. We defined the self-related brain regions as “self-referential” based on the Neurosynth meta-analysis map. Regions within the default mode network were based on Andrews-Hanna et al. (2010). For the self-related ROIs, we used all voxels within each cluster. For the ROIs from the default mode network, we used a 9-mm sphere surrounding the center coordinate (maximum of 123 voxels). *** $p < 0.001$, ** $p < 0.01$, and * $p < 0.05$ (uncorrected). All $p$ values rely on permutation test (1,000 times). *n.s.* non-significant. dmPFC, dorsomedial prefrontal cortex. PCC, posterior cingulate cortex. TPJ, temporoparietal junction. TempP, temporal pole. vmPFC, ventromedial prefrontal cortex. amPFC, anterior-medial prefrontal cortex. TC, temporal cortex. PHC, parahippocampal cortex. pIPL, posterior inferior parietal lobule. Rsp, retrosplenial cortex. TempP, temporal pole.

**Supplementary Table 1. Areas activated by the self > semantic contrast.**

| Location | MNI coordinates | | | Z | Cluster size (voxels) |
|---|---|---|---|---|---|
| | x | y | z | | |
| amPFC | 6 | 53 | 14 | 6.16 | 2,799 |
| vmPFC | -9 | 47 | -7 | 5.76 | |
| dmPFC | -9 | 56 | 26 | 5.48 | |
| SMA | -9 | 20 | 62 | 5.4 | |
| Left caudate | -12 | 8 | 11 | 4.42 | |
| Precuneus | -6 | -55 | 23 | 5.66 | 292 |
| Left insula | -33 | 20 | -16 | 5.15 | 175 |
| Left ANG | -51 | -70 | 32 | 4.5 | 138 |

amPFC anterior medial prefrontal cortex; vmPFC, ventral medial prefrontal cortex; dmPFC, dorsomedial prefrontal cortex; SMA, supplementary motor area; ANG, angular gyrus. A statistical threshold was set at $p < 0.001$ voxel-wise (uncorrected) and a cluster-level threshold was set a $p < 0.05$ (FWE corrected for multiple comparisons).

**Supplementary Table 2. Areas activated by the other > semantic contrast.**

| Location | MNI coordinates | | | Z | Cluster size (voxels) |
|---|---|---|---|---|---|
| | x | y | z | | |
| Precuneus | -9 | -55 | 20 | 6.66 | 487 |
| vmPFC | 3 | 50 | -13 | 6.16 | 1,518 |
| amPFC | 3 | 53 | 20 | 5.85 | |
| Rectus | 6 | 32 | -16 | 5.29 | |
| ACC | -3 | 32 | -1 | 5.14 | |
| Left MTG | -57 | -64 | 20 | 5.18 | 224 |
| Left ANG | -54 | -67 | 26 | 4.98 | |
| Right insula | 30 | 14 | -19 | 4.75 | 267 |
| Right MTG | 60 | 5 | -25 | 4.70 | |
| Left anterior MTG | -57 | -7 | -16 | 4.52 | 194 |

vmPFC, ventral medial prefrontal cortex; amPFC anterior medial prefrontal cortex; ACC, anterior cingulate cortex; MTG, middle temporal gyrus; ANG, angular gyrus. A statistical threshold was set at $p < 0.001$ voxel-wise (uncorrected) and a cluster-level threshold was set a $p < 0.05$ (FWE corrected for multiple comparisons).

**Supplementary Table 3. Areas activated by the introspection > categorization contrast.**

| Location | MNI coordinates | | | Z | Cluster size (voxels) |
|---|---|---|---|---|---|
| | x | y | z | | |
| dmPFC | -9 | 59 | 26 | 7.60 | 2,946 |
| vmPFC | -6 | 38 | -10 | 6.52 | |
| SMA | -6 | 17 | 62 | 6.25 | |
| Rectus | -6 | 41 | -19 | 6.18 | |
| MCC | -3 | -16 | 35 | 5.37 | |
| Left MFG | -27 | 44 | 20 | 4.64 | |
| Left temporal pole | -36 | 17 | -22 | 6.79 | 1,832 |
| Left insula | -30 | 14 | -19 | 6.65 | |
| Left IFG (orbital part) | -39 | 32 | -13 | 5.80 | |
| Left temporal pole | -45 | 14 | -28 | 5.42 | |
| Left ITG | -45 | 5 | -34 | 5.37 | |
| Left SMG | -57 | -43 | 26 | 5.02 | |
| Left MOG | -42 | -88 | 5 | 4.92 | |
| Right cerebellum | 30 | -79 | -34 | 6.18 | 204 |
| Right IOG | 45 | -82 | -1 | 6.12 | 1,258 |
| Right MOG | 42 | -85 | 2 | 5.97 | |
| Right IFG (orbital part) | 30 | 14 | -22 | 5.67 | |
| Right MTG | 48 | -34 | -4 | 5.46 | |
| Right temporal pole | 51 | 17 | -28 | 5.10 | |
| Right SOG | 21 | -97 | 11 | 4.73 | |

dmPFC, dorsomedial prefrontal cortex; vmPFC, ventral medial prefrontal cortex; SMA, supplementary motor area; MCC, middle cingulate cortex; MFG, middle frontal gyrus; IFG, inferior frontal gyrus; ITG, inferior temporal gyrus; SMG, superior marginal gyrus; MOG, middle occipital gyrus; IOG, inferior occipital gyrus; MTG, middle temporal gyrus; SOG, superior occipital gyrus. A statistical threshold was set at $p < 0.001$ voxel-wise (uncorrected) and a cluster-level threshold was set a $p < 0.05$ (FWE corrected for multiple comparisons).

**Supplementary Table 4. Areas activated by the memory > knowledge contrast.**

| Location | MNI coordinates | | | Z | Cluster size (voxels) |
|---|---|---|---|---|---|
| | x | y | z | | |
| Left MTG | -54 | -49 | 23 | 7.36 | 11,898 |
| MCC | -9 | -46 | 35 | 7.19 | |
| amPFC | 0 | 53 | 11 | 7.01 | |
| Precuneus | 9 | -49 | 20 | 6.70 | |
| Right IFG (orbital part) | 51 | 35 | -7 | 6.65 | |
| Left IFG (orbital part) | -36 | 20 | -16 | 6.50 | |
| Left cuneus | -12 | -58 | 23 | 6.47 | |
| Left insula | -30 | 17 | -16 | 6.42 | |
| Right IFG (opercular part) | 54 | 20 | 38 | 6.28 | |
| Left cerebellum | -21 | -79 | -40 | 6.18 | 427 |
| Right cerebellum | 18 | -73 | -34 | 5.32 | 215 |

MTG, middle temporal gyrus; MCC, middle cingulate cortex; amPFC, anterior medial prefrontal cortex; IFG, inferior frontal gyrus. A statistical threshold was set at $p < 0.001$ voxel-wise (uncorrected) and a cluster-level threshold was set a $p < 0.05$ (FWE corrected for multiple comparisons).

**Supplementary Table 5. RSA results: the Self = Other model.**

| Location | MNI coordinates | | | Z | Cluster size (voxels) |
|---|---|---|---|---|---|
| | x | y | z | | |
| Precuneus | -6 | -55 | 29 | 7.62 | 2,272 |
| MCC | 6 | -13 | 35 | 4.46 | |
| Right paracentral lobule | 12 | -34 | 50 | 4.30 | |
| amPFC | 3 | 50 | 14 | 7.50 | 16,577 |
| Right IFG (orbital part) | 45 | 44 | -4 | 6.33 | |
| Left ANG | -42 | -64 | 26 | 6.22 | |
| Right SFG (dorsolateral part) | 18 | 38 | 44 | 6.21 | |
| Left MFG | -24 | 32 | 41 | 5.97 | |
| Left SFG (dorsolateral part) | -18 | 35 | 47 | 5.93 | |
| Right IFG (triangular part) | 48 | 35 | -1 | 5.90 | |
| Left MTG | -60 | -7 | -19 | 5.82 | |
| Left IFG (orbital part) | -33 | 29 | -16 | 5.57 | |
| Left rectus | -12 | 38 | -16 | 5.38 | |
| Right cerebellum | 33 | -76 | -34 | 5.91 | 444 |

MCC, middle cingulate cortex; amPFC, anterior-medial prefrontal cortex; ACC, anterior cingulate cortex; IFG, inferior frontal gyrus; ANG, angular gyrus; SFG, superior frontal gyrus; MFG, middle frontal gyrus; MTG, middle temporal gyrus. A statistical threshold was set at $p < 0.005$ voxel-wise (uncorrected) and a cluster-level threshold was set a $p < 0.05$ (FWE corrected for multiple comparisons).

**Supplementary Table 6. RSA results: the Self = Introspection model.**

| Location | MNI coordinates | | | Z | Cluster size (voxels) |
|---|---|---|---|---|---|
| | x | y | z | | |
| dmPFC | 0 | 53 | 35 | 6.66 | 1, 930 |
| amPFC | 0 | 50 | 11 | 5.93 | |
| SMA | -12 | 20 | 62 | 5.81 | |
| Left MFG | -27 | 50 | 20 | 3.83 | |
| Left IFG (triangular part) | -51 | 20 | 2 | 4.77 | 379 |
| Left IFG (orbital part) | -45 | 20 | -10 | 4.10 | |
| Left ITG | -45 | 2 | -34 | 3.58 | |
| Left MTG | -57 | 8 | -28 | 3.13 | |
| Left temporal pole | -45 | 17 | -37 | 3.01 | |

dmPFC, dorsomedial prefrontal cortex; amPFC, anterior-medial prefrontal cortex; SMA, supplementary motor area; MFG, middle frontal gyrus; IFG, inferior frontal gyrus; ITG, inferior temporal gyrus; MTG, middle temporal gyrus. A statistical threshold was set at $p < 0.005$ voxel-wise (uncorrected) and a cluster-level threshold was set a $p < 0.05$ (FWE corrected for multiple comparisons).

**Supplementary Table 7. RSA results: the Self = Memory model.**

| Location | MNI coordinates | | | Z | Cluster size (voxels) |
|---|---|---|---|---|---|
| | x | y | z | | |
| Left IFG (orbital part) | -45 | 26 | -7 | 5.00 | 368 |
| amPFC | -12 | 44 | 5 | 3.48 | 103 |

IFG, inferior temporal gyrus; amPFC, anterior medial prefrontal cortex. A statistical threshold was set at $p < 0.005$ voxel-wise (uncorrected) and a cluster-level threshold was set a $p < 0.05$ (FWE corrected for multiple comparisons).

**Supplementary Table 8. Classifier-based MVPA results: Self>Semantic vs. Other>Semantic.**

| Location | MNI coordinates | | | Z | Cluster size (voxels) |
|---|---|---|---|---|---|
| | x | y | z | | |
| Right SFG | 18 | 35 | 38 | 5.69 | 4,214 |
| amPFC | 9 | 56 | 17 | 5.58 | |
| vmPFC | 6 | 50 | -10 | 5.06 | |
| Right MFG | 30 | 23 | 50 | 4.63 | |
| Rectus | 0 | 38 | -22 | 4.48 | |
| Right MFG (orbital part) | 24 | 47 | -16 | 4.40 | |
| Precuneus | 0 | -58 | 29 | 5.01 | 808 |
| MCC | 0 | -43 | 35 | 3.70 | |
| Calcarine sulcus | -3 | -70 | 11 | 2.89 | |
| Left MTG | -57 | -13 | -19 | 4.52 | 952 |
| Left IFG (orbital part) | -48 | 35 | -4 | 3.95 | |
| Left insula | -30 | 14 | -16 | 3.92 | |
| Left ITG | -54 | -7 | -31 | 3.91 | |
| Left IFG (triangular part) | -51 | 32 | 5 | 3.17 | |

SFG, superior frontal gyrus; amPFC, anterior medial prefrontal cortex; vmPFC, ventral medial prefrontal cortex; MFG, middle frontal gyrus; MCC, middle cingulate cortex; MTG, middle temporal gyrus; IFG, inferior frontal gyrus; ITG, inferior temporal gyrus. A statistical threshold was set at $p < 0.005$ voxel-wise (uncorrected) and a cluster-level threshold was set a $p < 0.05$ (FWE corrected for multiple comparisons).

**Supplementary Table 9. Classifier-based MVPA results: Self>Semantic vs. Introspection>Categorization.**

| Location | MNI coordinates | | | Z | Cluster size (voxels) |
|---|---|---|---|---|---|
| | x | y | z | | |
| Left MTG | -45 | -64 | 23 | 7.74 | 31,654 |
| dmPFC | 15 | 47 | 29 | 7.46 | |
| Precuneus | -6 | -58 | 14 | 7.39 | |
| Right MOG | 36 | -70 | 35 | 7.09 | |
| Left MFG | -21 | 41 | 32 | 6.96 | |
| PCC | 3 | -46 | 32 | 6.85 | |
| Right SFG (dorsolateral part) | 21 | 26 | 44 | 6.80 | |
| Left IPL | -36 | -73 | 44 | 6.69 | |
| Right MFG | 27 | 32 | 35 | 6.67 | |
| Left SFG | -21 | 17 | 47 | 6.38 | |
| amPFC | -3 | 50 | 11 | 4.74 | |
| vmPFC | 3 | 50 | -7 | 4.62 | |

MTG, middle temporal gyrus; MOG, middle occipital gyrus; MFG, middle frontal gyrus; PCC, posterior cingulate cortex; SFG, superior frontal gyrus; IPL, inferior parietal lobule; amPFC, anterior medial prefrontal cortex; vmPFC, ventromedial prefrontal cortex. A statistical threshold was set at $p < 0.005$ voxel-wise (uncorrected) and a cluster-level threshold was set a $p < 0.05$ (FWE corrected for multiple comparisons).

**Supplementary Table 10. Classifier-based MVPA results: Self>Semantic vs. Memory>Knowledge.**

| Location | MNI coordinates | | | Z | Cluster size (voxels) |
|---|---|---|---|---|---|
| | x | y | z | | |
| Left MOG | -33 | -61 | 35 | 7.51 | 33,057 |
| Right SFG (dorsolateral part) | 15 | 47 | 29 | 7.46 | |
| Right ANG | 45 | -49 | 32 | 7.36 | |
| Left ANG | -42 | -58 | 32 | 7.27 | |
| Left IPL | -36 | -58 | 47 | 6.91 | |
| Precuneus | -12 | -49 | 26 | 6.87 | |
| Right IPL | 39 | -49 | 50 | 6.76 | |
| Cuneus | -6 | -73 | 35 | 6.74 | |
| Left SFG (dorsolateral part) | -18 | 26 | 47 | 6.66 | |
| vmPFC | 3 | 41 | -7 | 4.58 | |
| ACC | -6 | 38 | 8 | 3.89 | |
| Rectus | 12 | 32 | -16 | 3.57 | |

MOG, middle occipital gyrus; SFG, superior frontal gyrus; ANG, angular gyrus; IPL, inferior parietal lobule; vmPFC, ventromedial prefrontal cortex; ACC, anterior cingulate cortex. A statistical threshold was set at $p < 0.005$ voxel-wise (uncorrected) and a cluster-level threshold was set a $p < 0.05$ (FWE corrected for multiple comparisons).
